# Modeling Bias Toward Binding Sites in PDB Structural Models

**DOI:** 10.1101/2024.12.14.628518

**Authors:** Stephanie A. Wankowicz

## Abstract

The protein data bank (PDB) is one of the richest databases in biology. The structural models deposited have provided insights into protein folds, relationships to evolution, energy functions of structures, and most recently, protein structure prediction, connecting sequence to structure. However, the X-ray crystallography (and cryo-EM) models deposited in the PDB are determined by a combination of refinement algorithms and manual modeling. The intervention of human modeling leads to the possibility that within a single structure, there can be differences in how well parts of a structure are modeled and/or fit the underlying experimental data. We identified that small molecule binding sites are more carefully modeled and better match the underlying experimental data than the rest of the protein structural model. This trend persisted irrespective of the structure’s resolution or its overall agreement with the experimental data. The variation of modeling has implications for how we interpret protein structural models and use structural models in explaining mechanisms, structural bioinformatics, simulations, docking, and structure prediction, especially when drawing conclusions about binding sites compared to the rest of the protein.

## Introduction

Today, most protein machine learning algorithms rely on the accuracy of existing structural models in the protein structural databank (PDB). While these models have been invaluable in advancing our understanding of biology, models in the PDB vary in the goodness of fit for experimental data^1^. Understanding how and why structural models differ in their fit to experimental data is critical for the interpretation, reliability, and uncertainty of conclusions drawn from downstream analyses, including structural bioinformatics, machine-learning algorithms, docking, or conclusions from single or groups of protein structural models.

Global metrics, such as the crystallographic R-factor^2^, are commonly used to determine the ‘goodness’ of a structural model ^3^. While R-factors indicate how well a model matches the observed structure factors, they are not always effective in discriminating major errors in a model^4^. Additional metrics, including geometry-based assessments, offer supplementary insights into model quality^5^, yet these, too, predominantly operate at a global level. To address this limitation, recent efforts have focused on developing residue-level metrics that evaluate how well individual regions of a structure fit the experimental data^6^. These developments enhance our ability to assess model accuracy across protein regions, shedding light on potential patterns or biases that could affect downstream applications. Since the full structure of a protein is essential for understanding its biological function^7^, any local differences in how well the model fits experimental data could lead to inaccurate or misleading biological interpretations.

Recognizing that macromolecular X-ray crystallography models are constructed through both manual and computational methods, we aimed to investigate whether biases systematically influence the alignment of different regions with experimental data.

Macromolecular X-ray crystallography models are created through an iterative process of manual modeling and refinement to achieve optimal agreement between the atomic model and the experimental data (structure factors). Manual modeling involves the modeler (i.e. human user) adjusting the atomic model to improve the fit to real space electron density^8^. The user aims to resolve ambiguities in the map by adjusting the structure by repositioning a portion of the molecule, including fitting ligands or other non-protein atoms, such as water molecules. This updated model is then fed through refinement software, optimizing the model’s fit with the experimental data in reciprocal space while maintaining plausible geometry, given user-defined parameters^9,10^. Refinement also updated the phases of the map, providing the user with an improved real-space electron density map to adjust the model manually. However, during the manual modeling step, due to the number of ambiguities between the map and model that could be fixed, there is a tendency to focus on modeling the ‘important’ parts of the structure, such as a binding or other functional site, while ignoring other parts of the protein structure. This process could introduce human bias into deposited structural models, leading to certain regions of the structure aligning more accurately with the experimental data than others.

Here, we explored whether there are local differences in how well parts of protein structures are modeled and determined how these differences likely came about. We observed that binding site residues frequently fit the experimental data better than those outside the binding site. Our findings suggest that this is primarily due to the more careful manual modeling of binding site residues, as evidenced by a higher number of alternative conformations and a better fit of geometry outliers to the experimental data. This suggests that these differences are likely attributable to user bias in generating structural models, potentially impacting conclusions drawn from individual structural models and influencing the performance of machine learning algorithms trained on these structures.

## Results

### Dataset

To create our initial dataset, we collected the fully optimized X-ray crystallography structures in the PDBRedo, a database of re-refined and re-built structures^11–13^(*n=68*,*332*). PDBRedo pipeline includes an automated re-refinement procedure with REFMAC^14^, rebuilding programs, including loop rebuilding and peptide flips^15,16^, and validation. More details can be found in Joosten et al^13^. This collection allowed us to focus on differences in manual modeling, which are not corrected in PDBRedo, rather than differences in refinement programs.

We first classified structures as bound or apo. To be classified as a bound structure, the structure ligand must have at least ten heavy atoms minus crystallographic additives (**Supplementary Table 1**). We then defined binding site residues as any residue within 5 angstroms (Å) of the largest ligand in the liganded structure. If the structure had multiple of the same ligand bound, due to symmetry or a natural homo-oligomer, all residues around each ligand were considered the binding site. These were defined as bound structures (*bound dataset; n=24*,*216*). We then obtained another dataset deemed ‘unbound’ structures with no ligand bound. These structures were allowed to have ligands in the crystallographic dataset (**Supplementary Table 1**) or ligands with less than 10 heavy atoms. To determine potential binding or functional sites in unbound structures, we used the fpocket algorithm^17^, which uses Voronoi tessellation and alpha spheres to identify pockets with structures. Pockets with volumes between 300 and 2000 Å^3^ were considered a possible binding site (*pocket dataset; n=21*,*381*). We then used the sphere representation of fpocket to determine all residues within the largest apo pocket, again using 5Å as the cut-off for pocket residues (**Supplementary Figure 1**). Other structures had either no pocket large enough or intermediate ligands.

### Density Fitness Calculations

We first aimed to determine if there were differences in the fit to electron density between binding site residues compared to all residues. To explore this, we obtained three metrics from PDBRedo that measure the fit of real space electron density to the model: real-space correlation coefficient (RSCC), real space R-values (RSR), and the mean electron density metrics for individual atoms (EDIAm) of residues^18,19^. RSCC measures the overall correlation between the observed density and the calculated density obtained from the model, with higher values indicating a better cross-correlation between the electron density and the model. RSR quantifies the deviation between the observed and calculated electron densities, with lower values indicating a better fit^4^. EDIAm evaluates the average electron density around each modeled atom, reflecting how well the electron density matches the expected value. Each of these metrics was determined on an individual residue basis, considering the main (including Cβ) and side chain atoms (**Methods**).

Overall, we observed that most residues fit the density well (*all residues, n=40*,*542*,*951*). However, with all real space electron density metrics, binding site residues fit the electron density better (*binding site residues, n=322*,*989*), **Figure 1A-C; Supplementary Table 2**). Across all residues, the average RSCC was 0.94 (Standard Deviation [SD]: 0.026), whereas binding site residues had an average RSCC of 0.96 (SD: 0.029), indicating an improved fit (**Figure 1A**; p=0.0, paired t-test). RSR also indicated a large difference in the fit to electron density between binding site residues and the entire protein, with an average of 0.058 (SD: 0.016) for binding site residues and 0.076 (SD: 0.026) for all residues (**Figure 1B**; p=0.0, paired t-test). While this metric again indicated that most residues fit well with the data, given it’s more sensitive to outliers than RSCC, it is likely identifying that there are more poor-fitting residues outside the binding site. EDIAm showed the same trend as RSR and RSCC (**Figure 1C**; binding site residues: 0.77 (SD: 0.198), non-binding site residues: 0.66 (SD: 0.162; p=0.0, paired t-test). While all metrics showed the same trend of binding site residues fitting to the experimental data better, it was consistent that there was a wider range of value in binding site residues, represented by higher standard deviations, potentially reflecting differences in modelers or due to other factors, such as resolution or global metrics of fit to data.

**Figure 1.**
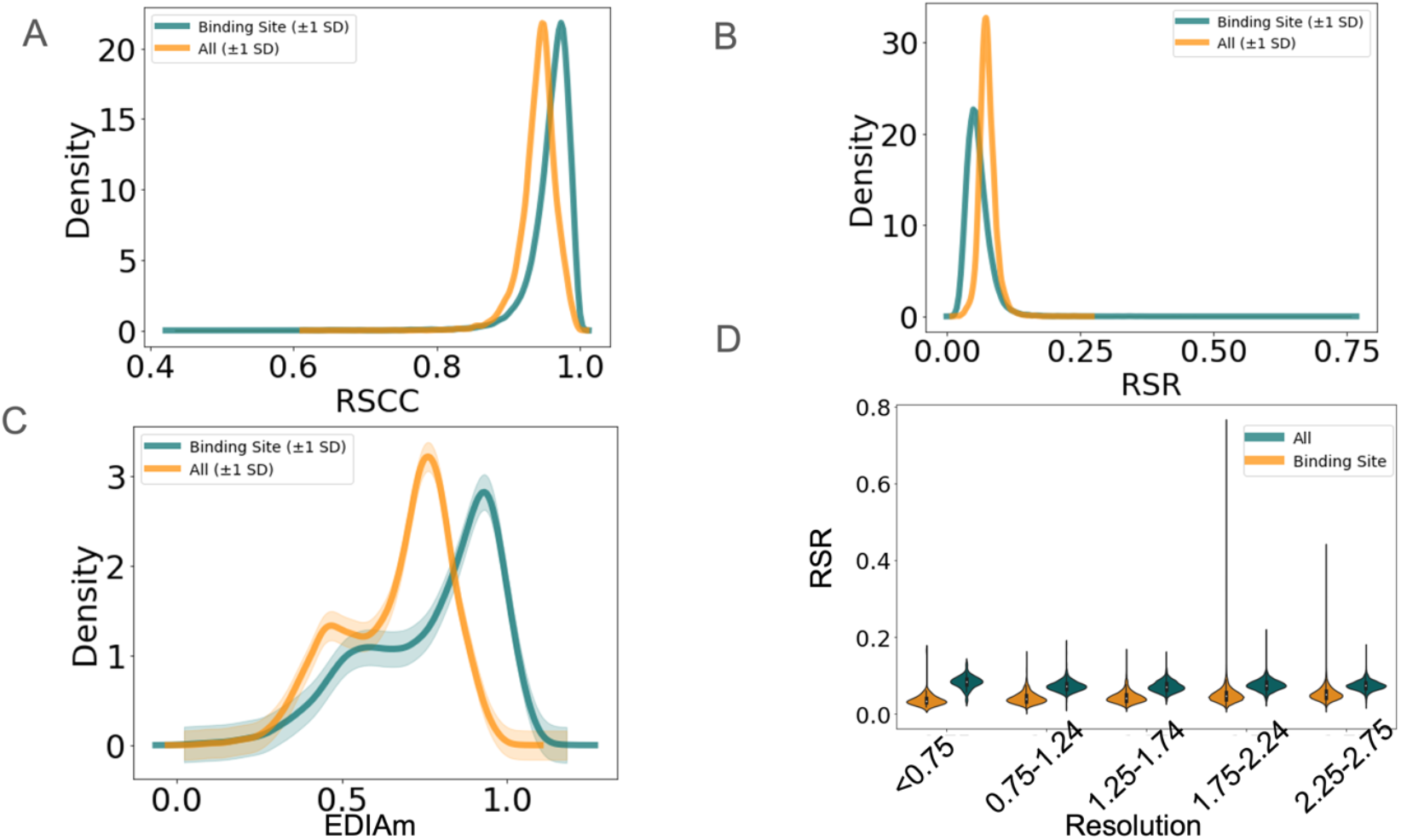
Binding site residues have a much better fit to real space electron density compared to non-binding site residues (**A**) RSCC metrics (binding site median-; non-binding site median; p=0.0), (**B**) RSR metrics (binding site median-; non-binding site median; p=0.0), (**C**) EDIAm metrics (binding site median-; non-binding site median; p=0.0), (**D**) RSR across resolution groups.

To explore how the fit to the electron density of the binding site versus all residues is impacted by resolution and global metrics, such as Rfree, we explored how the real space metrics differed by resolution and Rfree values. We observed that binding site residues fit the experimental data compared to all residues across all resolution ranges (0.5Å to 2.0Å; **Figure 1D; Supplementary Figure 2A**; Resolution: <0.75: Binding-mean 0.039, Non-binding-0.078; 0.75-1.24: Binding-0.044, Non-binding-0.074; 1.25-1.74: Binding-0.048, Non-binding-0.074; 1.75-2.24: Binding-0.054, Non-binding-0.076). However, these trends were more pronounced in higher-resolution structures, indicating that selecting structures based on resolution would not eliminate the bias of fit to density. We also observed that the trend of binding site residues having a better fit to the electron density holds across a range of Rfree values. Similar to resolution, the trend is more pronounced with lower Rfree. The smaller differences between binding site residues and all residues in lower resolution and higher Rfree structures are likely due to increased poor fitting of all residues. Together, these results indicate that global metrics of data fit also do not eliminate this bias (**Supplementary Figure 2B/C**).

Based on these results, we wanted to determine what might be driving the differences between binding site residues and the entire structure. One possibility is structural, where binding site residues may exhibit reduced solvent exposure, increased rigidity, or other structurally significant characteristics. To explore this, we examined possible binding site pockets containing no ligand (*pocket dataset*). This set of pocket residues should have similar underlying structures to residues around binding site residues. While the fit of pocket residues (EDIAm: 0.77, SD: 0.151) to the underlying electron density, measured with EDIAm was better than all residues (0.66, SD: 0.143), there was a much smaller, but still statistically significant difference (paired t-test: 0.0, **Supplementary Figure 2D**).

Given the smaller differences in pocket residues, along with variations in resolutions, R-free values, and the large standard deviation in the fit of binding site residues to experimental data, this suggests that the observed distribution of differences in the fit may stem from manual modeling variations. This prompts us to investigate how the binding site residues were modeled in comparison to the rest of the protein.

### Number of Alternative Conformations

To determine what was driving these differences in fit to density, we first explored if it was driven by the number of alternative conformations (altlocs) placed. Crystallographers utilize altlocs to describe parts of the structure that could reasonably be modeled in multiple discrete locations. While there are now multiple algorithms to place alternative conformations automatically^20–22^ and help guide users in Coot where alternative conformations should be placed^23^, most alternative conformations are placed manually. Therefore, we took this as a metric of modelers paying closer attention to areas of the protein with more alternative conformations.

We observed that over 58% of structures have at least one altloc. In binding site residues, 5.0% (SD: 0.13) had at least one altloc compared to 1.9% (SD: 0.06) of residues elsewhere in the protein (**Figure 2A;** p=5.07e-165, paired t-test). We interpreted this finding as modelers paying more attention to binding site residues than the rest of the protein. Given that altlocs are more likely to be placed at high resolution and in models with better global metrics of fit, we explored if these trends changed across different resolution ranges and/or Rfree values (**Figure 2B; Supplementary Figure 3A**). Similar to electron density metrics, differences in the number of altlocs placed were most pronounced in high-resolution structures (resolution better than 1.74Å). As altlocs are more frequently placed in higher-resolution structures, it is unsurprising that lower-resolution structures have a less pronounced difference in the number of altlocs placed.

**Figure 2.**
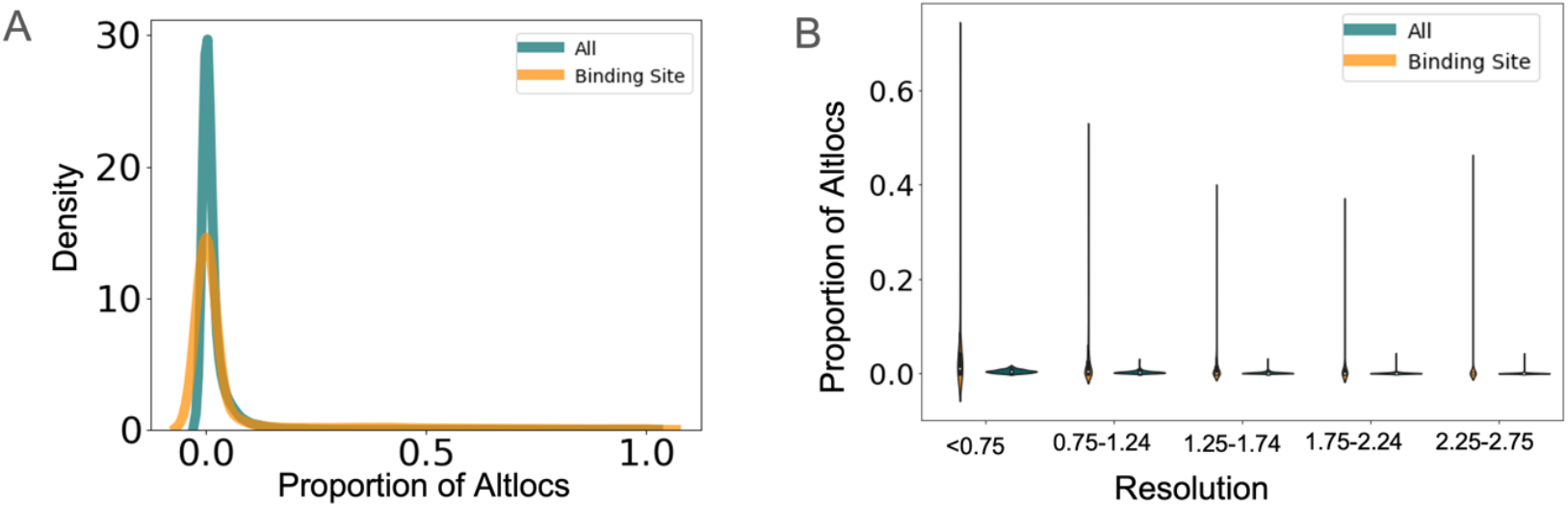
(**A**) Altlocs are more frequently placed in binding site residues compared to all residues. (**B**) Altlocs are more frequently placed in binding sites, regardless of the resolution of the structure.

We then explored how many altlocs are placed in pocket residues versus the rest of the protein. While there was an increase in the number of altlocs in pocket residues compared to all residues, this difference was much less pronounced than in binding site residues around a small molecule or other binding event (**Supplementary Figure 3B**), similar to what was observed with the fit to electron density. This indicated that the results we observed around ligand binding sites were likely due to modeling differences rather than biological differences.

### Geometry Metrics

We then explored if the differences in fit to density might be driven by improved fitting of individual residues, regardless of the number of conformations. To explore this, we looked at side chain geometry metrics. Beyond placing ligands and/or altlocs in protein structures, side-chain rotamers are often manually altered^8^. While refinement programs restrain structures based on idealized geometric metrics^10,24^, including fitting into idealized rotamer space, these can be changed manually. While there are cases, especially in high-resolution structures, that rotamers should fit outside of their ‘idealized’ position^25^, we expected that there would be fewer outliers in a more carefully modeled area of the protein, such as binding site residues. To explore if the geometry is better in binding site residues, we examined the rotamer status of every residue, examining how many were labeled as ‘outliers’ or ‘allowed’ from Phenix.rotalyze^26^. Phenix.rotalyze classifies rotamers as outliers if their occurrence in the reference dataset^27^ is below 0.3%, while rotamers categorized as allowed have an occurrence between 0.3% and 2.0% in the reference data^26^. In binding site residues, 2.4% were classified as outliers, compared to 2.0% across all residues (**Figure 3A**; p=0.56). We then examined the allowed rotamers, finding that 4.2% of binding site residues were classified as allowed, compared to 4.8% of residues across the entire dataset (**Figure 3B;** p=1.38e-08).

**Figure 3.**
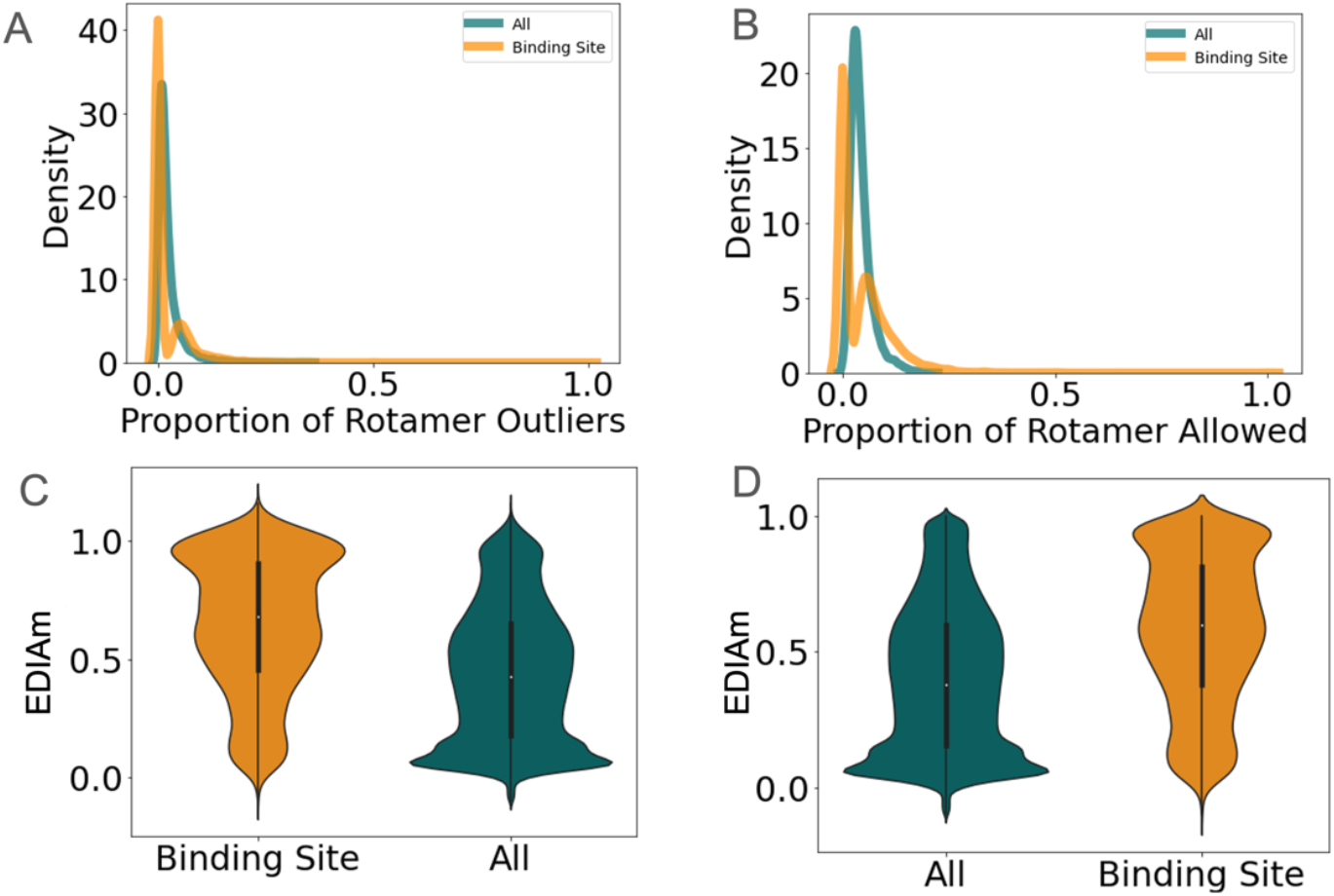
Binding site residues have fewer non-favored rotamer values. (**A**) Proportion of residues with rotamer outliers (binding site-0.009; all residues-0.015; p=8.79e-42), (**B**) Proportion of residues with allowed rotamer (binding site-0.42; all residues-0.48; p= 8.79e-42) (**C**) Distribution of EDIAm in residues classified as rotamer outliers (Binding site-0.52; All residues-0.36; p=1.3e-40) (**D**) Distribution of EDIAm in residues classified as allowed rotamer (binding site-0.64; all residues-0.43; p=5.89e-95)

As some rotamer outliers represent true unusual positioning of residues^25^, especially when interacting with a substrate or small molecules, we suspected that this was driving the bimodal distribution only observed in binding site residues. We hypothesized that binding site residues with rotamer outliers would have better evidence from the electron density to support their outlier status. Therefore, we explored whether outlier rotamers in the binding site had better evidence from the experimental data, comparing the EDIAm values of outliers found in binding sites versus all residues. We observed that binding site residues classified as outliers (**Figure 3C**) and those modeled as allowed (**Figure 3D**) had an improved fit to the experimental data compared to all residues with the same classifications. This suggests that non-ideal rotamers in binding sites are more likely to result from genuine geometry outliers, supported by density data, compared to residues with non-ideal rotamers outside of the binding site. Together, this supports the notion that users tend to model binding site residues more carefully, adjusting their geometry to deviate from the ‘normal’ distribution when experimental data supports it. In contrast, residues outside the binding site likely undergo less meticulous manual modeling.

## Discussion

Over multiple metrics, we observed that binding site residues tend to fit experimental data more accurately. Our data indicates that this is likely driven by more carefully manually modeled binding site residues, as observed by increased number of alternative conformations, the reduction in geometry outliers, and the fit of geometry outliers to the underlying experimental data in binding site residues. Overall, this highlights a strong bias in the accuracy of structural models toward binding sites, with implications for drawing conclusions across studies from a single structural model to machine learning algorithms. We call on the structural biology community to try to reduce this bias modeling moving forward by paying attention to areas outside of the binding site and for the need for more automated or guided modeling algorithms^19–21,23^. Although manual modeling will likely remain a key component of experimental structural biology, the shift towards higher-throughput techniques and the increasing complexity of large cryo-EM complexes necessitate more automated experimental modeling methods, as manually handling larger models is becoming progressively more challenging.

While we focused our analysis on PDBRedo to eliminate errors due to varied refinement, this trend is likely worse within non-PDBRedo structures. PDBRedo has provided individual residue metrics, and these metrics should be used to highlight local errors in models. However, echoing what has been called for before^4^, within the PDB there is a pressing need for more useful measures to highlight local errors in structural models both upon deposition and for people who are downloading and using the data. These metrics are likely even more important for other frequently mismodeled molecules, such as ligands and waters^39^.

These biased structural models often serve as ‘truth’ for simulations, docking, bioinformatic studies, and, most recently, machine learning algorithms^24–30^. However, our findings emphasize that we must be careful in using and explaining biological results from PDB models. *It is critical to understand that PDB models are just that*, <u>***models***</u>. They do not fully explain the underlying experimental data, have nuances in how they are representing the data, including alternative conformations, anisotropic B-factors, or translation-liberation-screw parameters, and as we show, can vary across the model in how well the underlying data is explained due to user error or biases^31–33^. While there are efforts to harmonize and improve the encoding of this data to capture more of these nuances^34^, many structural biology machine learning algorithms directly use these biased models to draw biological conclusions, potentially perpetuating errors. The modeling biases that we present here, adds to additional biases in the PDB based on the ability to capture states of a macromolecule using X-ray crystallography or cryoEM^35^. Further, we and others have demonstrated that the structures and dynamics outside the binding site are important! These areas are critical for allostery, binding, and cooperativity, to name a few. Belittling these areas in our structural models can perpetuate biological conclusions about the importance of binding sites over the rest of the protein^10,36–38^. Understanding the errors, biases, and uncertainties in data is as important as the computational algorithms when drawing conclusions.

## Methods

### PDBRedo Database

The current entries of final optimized PDBs from the PDBRedo database (https://pdb-redo.eu/download) was download in November 2024. Structures that had a final pdb and corresponding mtz.

### Electron Density Fit Metrics

Metrics on the fit of electron density metrics (RSR, RSCC, EDIAm) were obtained from PDBRedo in the PDB_final.json files. Each of these metrics was determined on an individual residue basis, considering the main (including Cβ) and side chain atoms. More details can be found in references 18 and 19. The code to obtain these metrics is located at https://github.com/PDB-REDO/density-fitness.

### Alternative Conformations

To determine the percent of alternative conformations in each residue, we classified residues as having zero or more than one alternative conformation. If any heavy atom in the residue had an altloc, then the residue was considered to have an alternative conformation. We then determined the proportion of residues classified as having an alternative conformation.

### Rotamer

We used phenix.rotalyze (Phenix version 1.21) to determine each sidechain’s allowed or outlier status ^24^. Residues are categorized as favored, outliers, or disallowed based on their χ angles and how closely they match known rotameric states within the Dunbrack Rotamer Library^27^.

### fpocket

Fpocket was run with default settings (version 4.2.2; https://github.com/Discngine/fpocket), only specifying the file name. We then took the largest pocket identified and determined if the volume was between 300 and 2000 Å^3^.

---

**Supplementary Figure 1.**
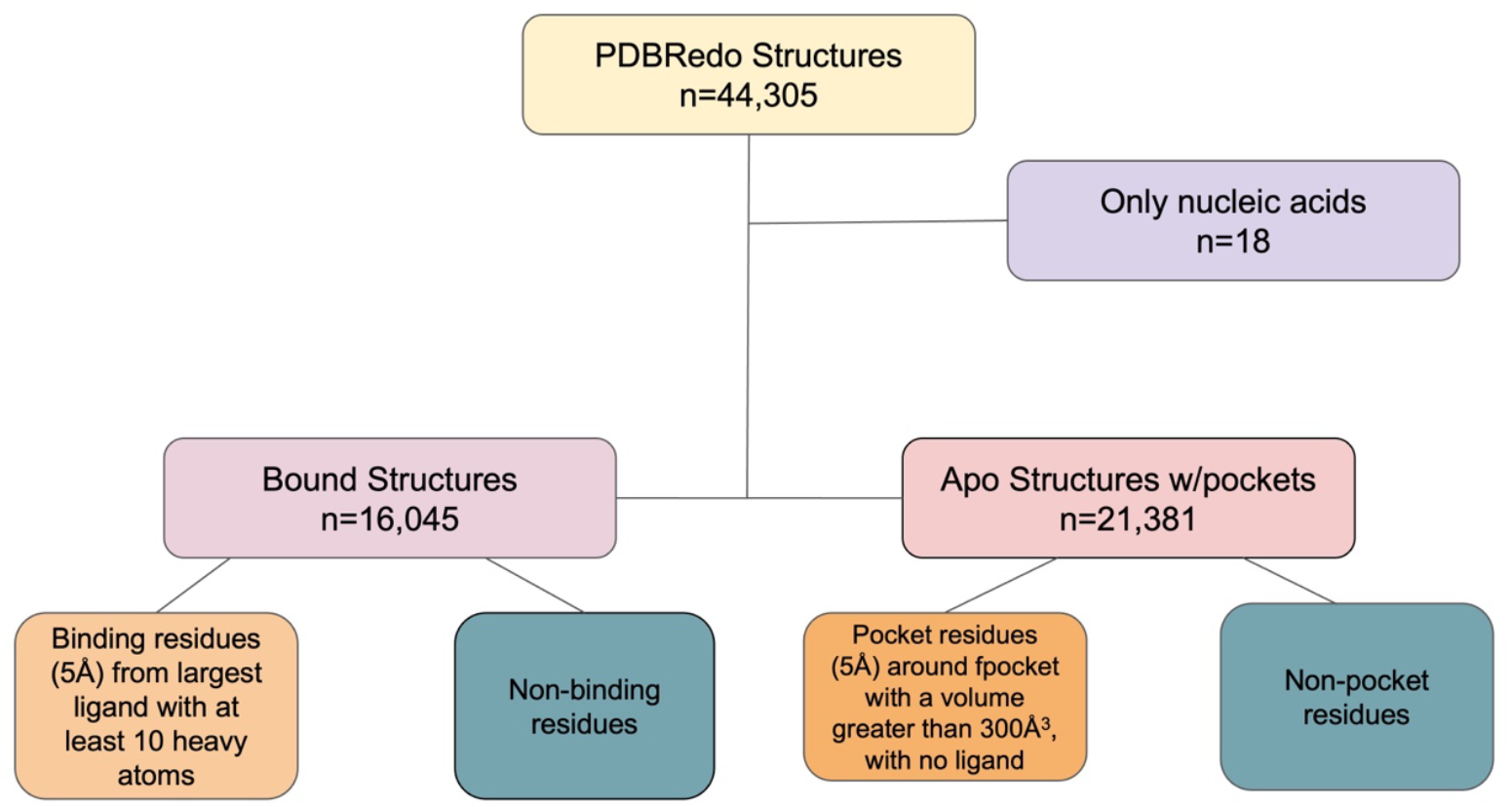
Workflow for determining PDBs and residues in comparison groups

**Supplementary Figure 2.**
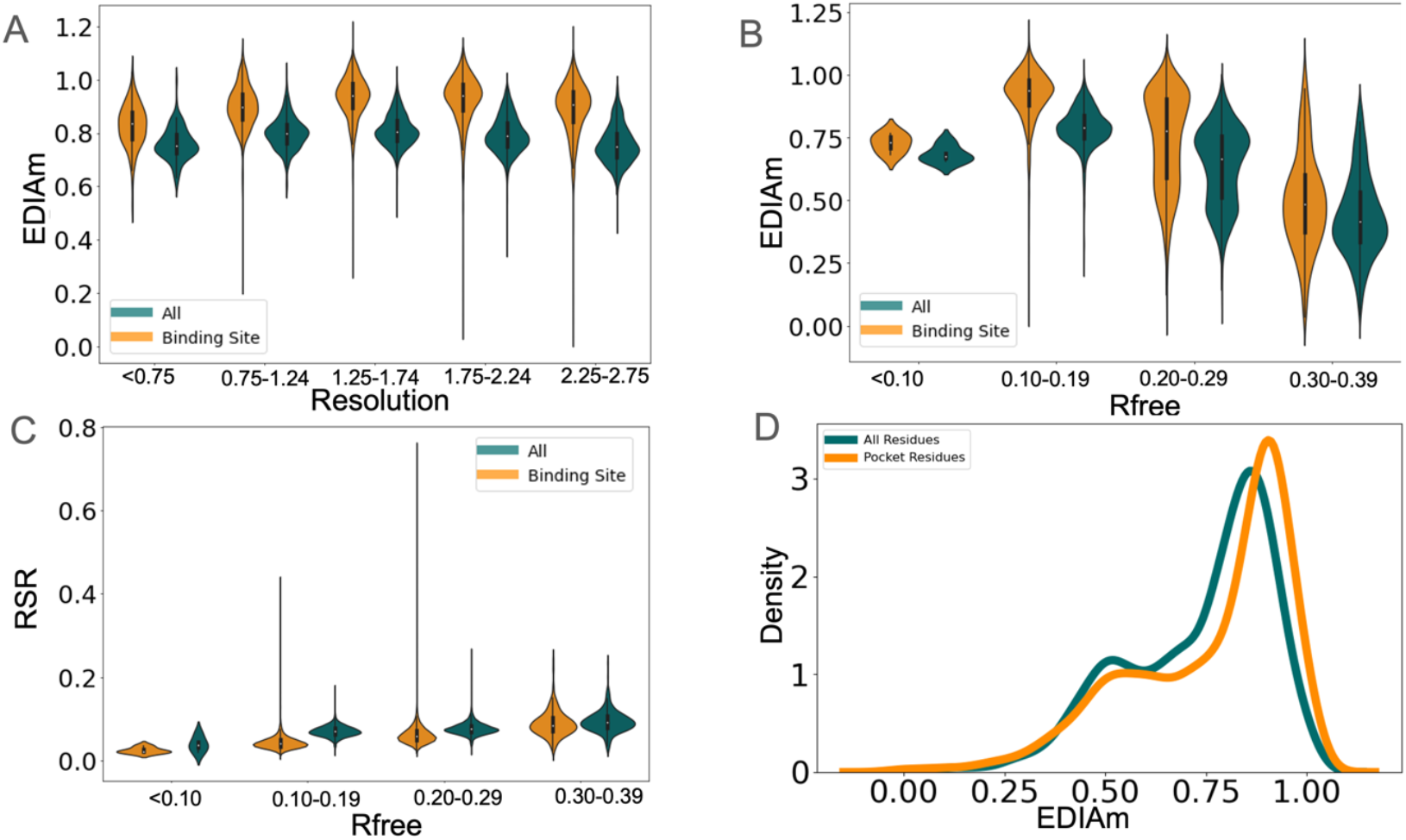
(**A**) EDIAm differences by resolution. While all groups were statistically significant, the largest difference were in the higher resolution groups. (**B**) EDIAm differences by Rfree value. We observed similar trend across all Rfree ranges, with Rfree between 0.10 and 0.19 having the most extreme differences (binding-0.89, all-0.79). (**C**) RSR differences by Rfree values. (D) EDIAm values in pocket residues versus all residues (pocket-0.72, all-0.66).

**Supplementary Figure 3.**
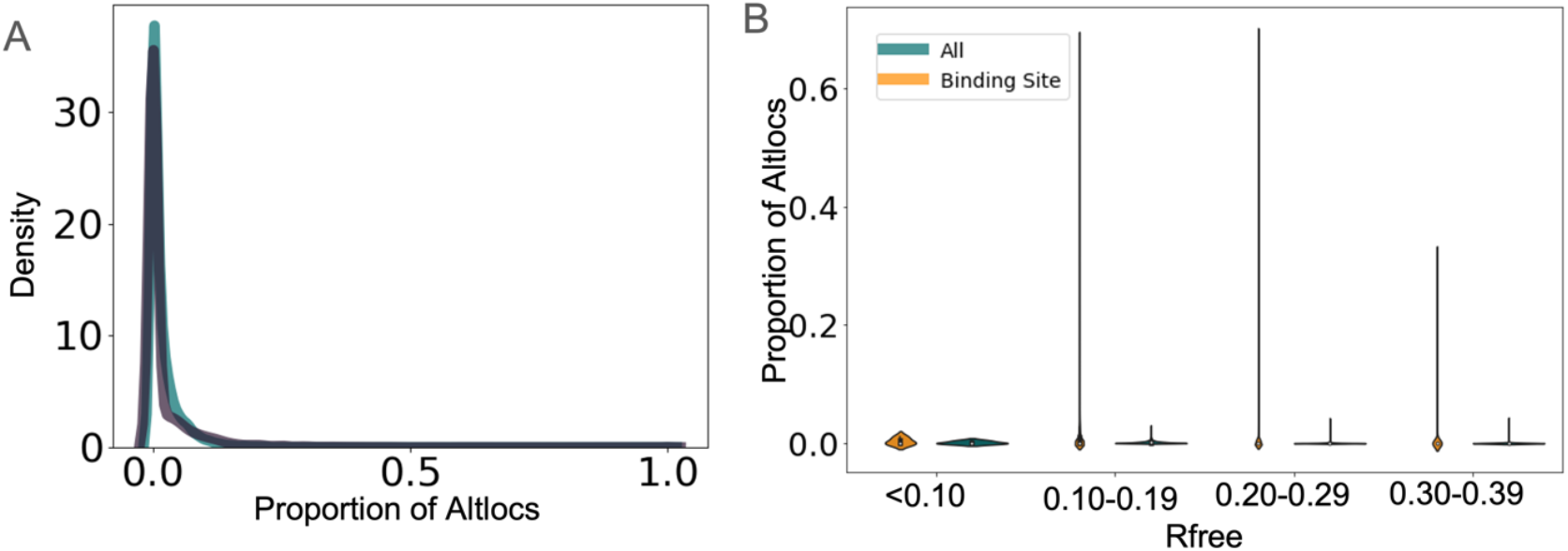
(**A**) The number of altlocs places in pockets without ligands occurs more often than non-pocket residues, but less frequently than in binding site residues (Pocket residues median-2.0%; all residues median-1.9%; p=3.38e-07). (**B**) Increased number of altlocs modeled in binding sites occurs regardless of Rfree values.

